# Mitotic Outcomes in Fibrous Environments

**DOI:** 10.1101/2021.11.21.469470

**Authors:** Aniket Jana, Haonan Zhang, Ji Wang, Rakesh Kapania, Nir Gov, Jennifer DeLuca, Amrinder S. Nain

**Affiliations:** Department of Mechanical Engineering, Virginia Tech, Blacksburg, VA 24061, USA; Department of Biomedical Engineering and Mechanics, Virginia Tech, Blacksburg, VA 24061, USA; Department of Aerospace and Ocean Engineering, Virginia Tech, Blacksburg, VA 24061, USA; Department of Chemical and Biological Physics, Weizmann Institute of Science, Rehovot, 7610001, Israel; Department of Biochemistry and Molecular Biology, Colorado State University, Fort Collins, CO 80523, USA

**Keywords:** mitotic cell rounding, mitotic forces, nanofibers, mitotic spindle, retraction fibers

## Abstract

During mitosis, cells round up and generate outward forces to create space and orient the mitotic spindles. Here, using suspended ECM-mimicking nanofiber networks, we recapitulate *in vivo* adhesion organization and confinement to interrogate mitotic outcomes for various interphase cell shapes. Elongated cells attached to single fibers through two focal adhesion clusters (FACs) at their extremities result in perfect spherical mitotic cell bodies that undergo large 3D displacement while being held by retraction fibers. Increasing the number of parallel fibers increases cellular extremity FACs and retraction fiber-driven stability, leading to reduced 3D cell-body movement, metaphase plate rotations, and significantly faster division times. Interestingly, interphase kite shapes on a crosshatch pattern of four fibers undergo mitosis resembling single-fiber outcomes due to rounded bodies being primarily held in position by retraction fibers from two perpendicular suspended fibers. We develop a cortex-astral microtubule analytical friction and force model to capture retraction-fiber-driven stability of the metaphase plate rotations. We report that reduced orientational stability results in increased monopolar mitotic defects. In the case of cells attached to two parallel fibers, rounded mitotic cells can get confined between the suspended fibers, allowing estimation of the mitotic forces through measurement of the outward deflection of the fibers. Interestingly, confinement causes rotated mitotic spindles similar to those reported in dense tissues. Overall, we establish dynamics of mitosis in fibrous environments governed by fiber arrangement and architecture-driven differences in interphase cell shapes, adhesion geometries, and varying levels of mechanical confinement.

## Introduction

The mitotic spindle is a remarkable biological machine responsible for the proper segregation of chromosomes during cell division^1,2^. Its positioning determines the cell division axis orientation and is thereby crucial for tissue morphogenesis. Defects in spindle positioning are often implicated in developmental disorders and cancer progression^3^. Mitotic spindle positioning depends upon several factors, including delamination^4^, cell shape^5^, the spatial organization of cell-matrix focal adhesions (FA) present during interphase^6–8^, and confinement forces.^9,10^ *In vivo* cells undergoing mitosis push on neighboring interphase cells to generate space for rounded cell shape for proper mitotic spindle positioning^11,12^. Rounded cells are held in place within tissues through basolateral septate junctions^4^, actin *mitotic* protrusions in 3D gels^13^, or the *in vitro* substrate via thin actin-rich structures, referred to as retraction fibers originating from active integrin sites^10,14,15^. *Mitotic* protrusions and retraction fibers act as mechanical links to stabilize the orientation of the mitotic spindle^6,16,10^ and to facilitate the spreading of daughter cells following cytokinesis through the re-establishment of FAs^14^.

Cells in the body can move along or between interstitial fiber networks composed of a loose network of fibers^17^. Very little is known about how mitosis proceeds in loose fibrous extracellular (ECM) environments. Can the organization of ECM fiber networks direct mitotic outcomes? Central to this inquiry is the finding that cells in native fibrous 3-dimensional environments form elongated cell-matrix adhesions at their peripheries that differ significantly from the low aspect ratio adhesions formed isotropically on flat 2D surfaces^18–24^. Mitosis on flat 2D and on 2D adhesive micropatterns designed to control cell shapes occurs primarily in an unrestricted manner^25,26^, while natural ECM imposes 3D-mechanical confinement^13,27,28^. Thus, we inquired if mitotic progression and outcomes differed in suspended fibers that mimic the natural interstitial ECM^29^.

Fibrous ECM consists of individual fibrils and bundled fibers ranging in sizes from a few hundreds of nm to several microns^30,31,32^, organized in a diverse range of fiber densities and network architectures, including aligned configurations^33^ and crossing fiber arrangements ^34–37^. We used our previously reported non-electrospinning ‘Spinneret Tunable Engineered Parameters’ (STEP) technique to generate aligned and crosshatch networks of suspended fibers ^38,39^. Controlling the interfiber spacing allowed us to achieve cells in elongated and kite shapes^19,20^, with varying density of focal adhesion clustering (two, four, and multi) at the cell peripheries. Retraction fibers originating from the focal adhesions held rounded cells in position but with the freedom to move about the fiber axes due to the absence of a basal surface, a key difference with 2D culture methods. Increased retraction fiber coverage induced higher stability, manifested in reduced cell body movement, metaphase plate oscillations, mitotic times, and a shift of mitotic errors from monopolar to multipolar spindle defects. A theoretical cortex-astral microtubule friction model developed captures retraction-fiber-driven stability of the metaphase plate rotations. Confinement of cells between two parallel fibers overrode retraction-fiber stability cues and unexpectedly caused a tilt of mitotic spindle, similar to those reported in dense tissue environments^4,40,41^. The tilt of mitotic spindles was discovered to be sensitive to the cellular position between the confining fibers, and using nanonet force microscopy (NFM)^42–44^, we estimated the outward forces exerted by cells during division. Overall, we present a new biologically relevant model system of suspended fiber networks that details mitotic rules in fibrous microenvironments mimicking the loose-interstitial tissue and dense tissues.

## Results

### Focal adhesion clustering controls metaphase plate dynamic repositioning

We wanted to inquire if the focal adhesion patterns on fibers correlating with different cell shapes and aspect ratios (AR) contributed to the dynamic orientation of the metaphase plate. We generated suspended nanofiber (fiber diameter ~ 250 nm) networks in both aligned and crosshatch geometries to achieve control on cells of diverse AR and FA clusters (**Fig. 1a**). Tuning the inter-fiber spacing (4 μm −25 μm) in the aligned networks resulted in 3 elongated high AR cell shapes, i) one fiber elongated (1F-elongated) shapes of cells attached to single fibers and having two major FA clusters, ii) two fiber-rectangular (2F-elongated) shaped cells attached to two fibers and having four FA clusters, and iii) multifiber-rectangular (MF-elongated) shaped cells attached to ≥ three fibers with multiple aligned FA clusters. Crosshatch networks of inter-fiber spacing ~ 50 μm induced symmetric polygonal kite-shaped cells with four FA clusters. Quantification of the ARs during interphase revealed the highest elongated geometries in the 1F-elongated and 2F-elongated categories (AR~9, **Supplementary Fig. S1**), followed by the MF-elongated shapes (AR ~ 5) and finally the 2F-kite shape and 2D cells on flat 2D having AR close to 1. Using HeLa cells expressing Histone H2B GFP, we investigated the dynamics of mitotic progression (Fig. 1a, and **Supplementary Movies 1-5**). As cells rounded up during mitosis, we observed them held in place by actin retraction fibers originating at the FA cluster sites and connecting to the cell cortex. We characterized the 3D shape of mitotic cells in this rounded state, specifically during metaphase. Quantification of the mitotic cell heights (**Fig. 1b**) revealed that cells adhering to suspended fibers can round up significantly more (increased cell height) than their counterparts in flat 2D. In flat surfaces, mitotic cells are not perfectly spherical but instead show an elliptical shape in their cross-sectional views (Fig. 1b), with aspect ratios (cell height by width) significantly less than 1 (0.84±0.09, Mean±SD, Fig 1b). In the 1F-elongated shape, we observed almost perfectly spherical mitotic cells (Fig 1b) as supported by our aspect ratio (1.02 ± 0.03, Fig 1b) and circularity (0.980 ± 0.014, Fig 1b) measurements. Interestingly, in the multifiber category, where cells are positioned on top of the fiber layer, the mitotic cell shape is significantly more flattened (Aspect ratio: 0.908 ± 0.077, Fig 1b) and also features cortical deformations (Fig 1b), thereby leading to a significant reduction in their circularity (0.915 ± 0.066, Fig 1b). In these respects, the MF system is more similar to cells on flat 2D surfaces.

**Figure 1:**
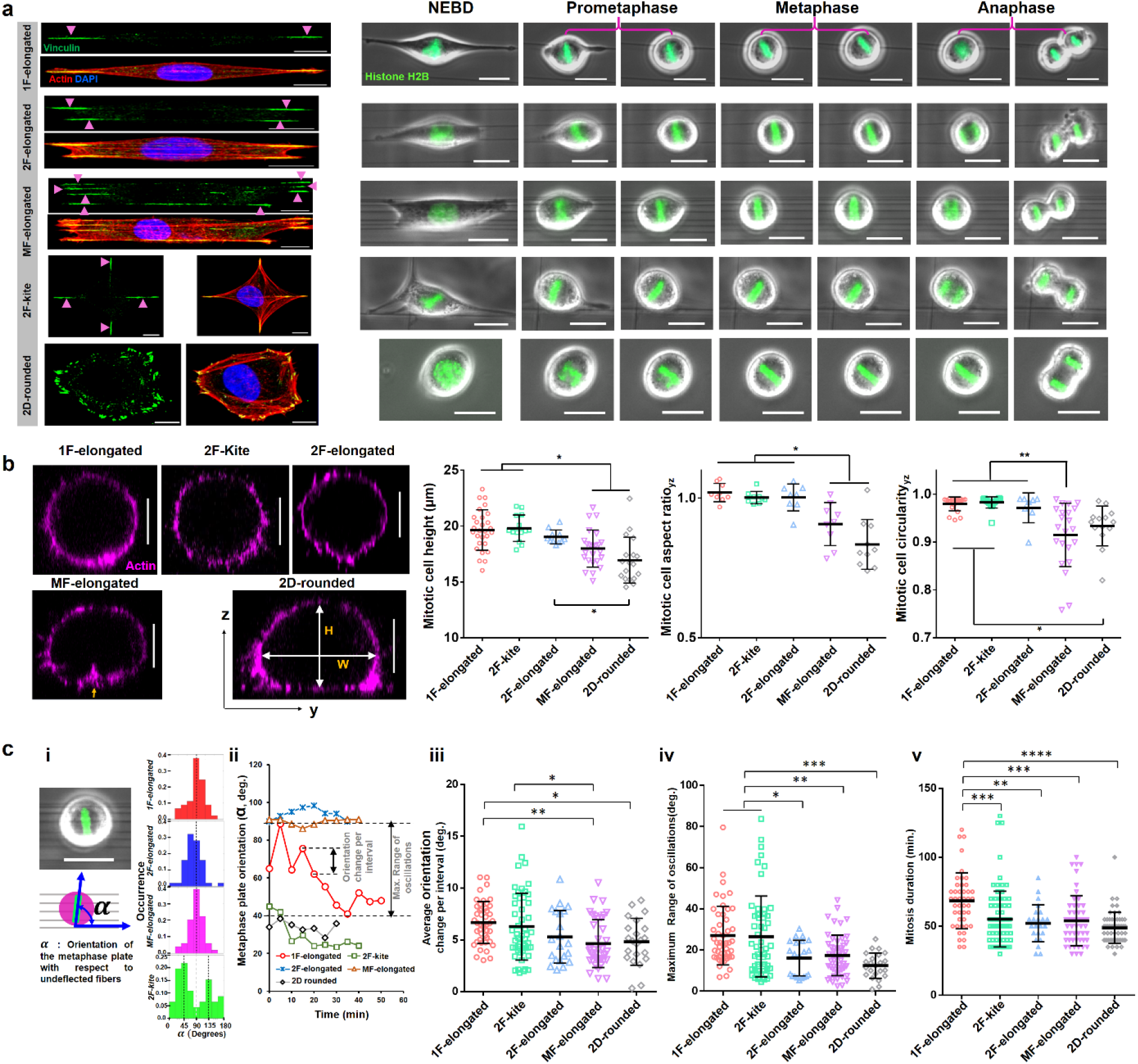
Cell division in suspended fiber environments. (a) Time-lapse images showing representative cases of mitotic progression, corresponding to 5 distinct interphase cell shapes: needle(1-fiber), 2F-rectangular (2 fibers), MF-rectangular (≥3 fibers), 2F-kite (2 orthogonal fibers) and 2D-rounded (glass coverslips). Cells are fluorescently stained with actin (red), vinculin (green) and nucleus (blue), Scale bars represent 10 μm and 20 μm respectively, (b) Representative yz cross-sectional images of actin stained cells fixed at metaphase on different substrate categories, mitotic cell height (H), mitotic cell aspect ratio (H/W) and mitotic cell circularity (4*π*Area/Perimeter2) for different substrate categories, (c,i) Illustration showing convention for metaphase plate orientation (α), (ii) Analysis of individual temporal profiles of metaphase plate orientation using orientation change per interval maximum range of metaphase plate oscillations (iii,iv) Statistical comparison across all fiber configurations and 2D for metaphase plate dynamics, (v) mitosis duration analysis showing 1F-elongated to be slowest.

Consistent with previously reported literature^45^, we observed that the metaphase plate (MP) exhibited significant fluctuations about its mean position following mitotic entry. We characterized the fluctuations of the MP orientation (*α*, **Fig. 1c(i)**, and **Supplementary Movies 6-9**) and found that in aligned fiber networks, the orientation profiles were centered around 90° (i.e. orthogonal to the cell elongation during interphase, **Fig. 1c(i)** and **Fig. S1**) in good agreement with Hertwig’s rule^46^. Interestingly, in the case of 2F-kite shaped cells, the temporal profiles (**Supplementary Fig. S2**) were mainly centered around 45° or 135°, indicating a diagonal placement of MP between the two orthogonal adhesion sites.

A closer inspection of the individual temporal profiles (**Fig. 1c(ii)**) indicated significant movement of MPs during mitosis. We quantified the degree of orientation changes using two metrics: i) absolute orientation change of the MP per time interval and ii) maximum angular range of MP rotations. Across all elongated shapes, we observed that the 1F-elongated shaped cells with two FA clusters demonstrated significantly higher orientation changes (**Fig. 1c(iii)**) and the highest angular range of MP rotations (**Fig. 1c(iv)**). Incidentally, similar dynamic activity levels of the MP were also observed in symmetric 2F-kite shaped cells with four FA clusters. Intrigued by the differences in MP dynamics and the curious similarity between 1F-elongated and 2F-kite shapes, next, we investigated the durations for mitosis completion (nuclear envelope breakdown to anaphase completion, **Fig. 1c(v)**). We found that 1F-elongated cells took significantly longer average times (68.4±3 min, mean±sem) to divide, as compared to other suspended cell shapes (50-55 min) or on traditional flat glass coverslips (48.8±1.5 min).

### The spatial organization of retraction fibers modulates mitotic cell shape and dynamics

Our observations of increased metaphase plate reorientations and mitotic times in 1F-elongated cells prompted us to inquire about the role of retraction fiber-driven stability in mitotic outcomes. We observed rounded cells held in position by retraction fibers (schematically shown in **Fig. 2a(i)**). Using confocal microscopy, we analyzed the TOP (xy, **Fig. 2a(ii)**), SIDE (zy, **Fig. 2b**), and FRONT (xz, **Fig. 2c(i)**) views. We identified retraction fibers to appear as actin-rich hotspots in the averaged intensity heatmaps of the cell cortex. 1F-elongated cells attached to single fibers, through two major FA clusters, were connected by two major sets of retraction fibers during mitosis. Interestingly, 2F-kite shaped cells, despite having four FA clusters during interphase, were found to be anchored by two dominant sets of orthogonally arranged retraction fibers in the rounded mitotic state. To quantitate the spatial localization of retraction fibers, we introduced a new metric: retraction fiber coverage (RFC, 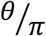), defined as the fraction of the cortical perimeter to which retraction fibers were connected, along two principal equatorial planes (**Fig. 2a(ii)** for TOP view, and **Fig. 2c(iv)** for FRONT view). Averaged over both planes, the 1F-elongated cells had an RFC of only ~0.19 and 0.15, while cells on Flat 2D had an average value of ~ 0.92 and 0.25 for TOP and FRONT views, respectively. Not surprisingly, plotting MP orientation changes per interval and maximum angular range, as a function of RFC, demonstrated reduced MP oscillations with increasing retraction fiber coverages (**Fig. 2a(v)**). Coincidentally, mitosis time decreased with an increasing number of retraction fibers (**Fig. 2a(vi)**), indicating a role of mechanical stability in mitosis.

**Figure 2.**
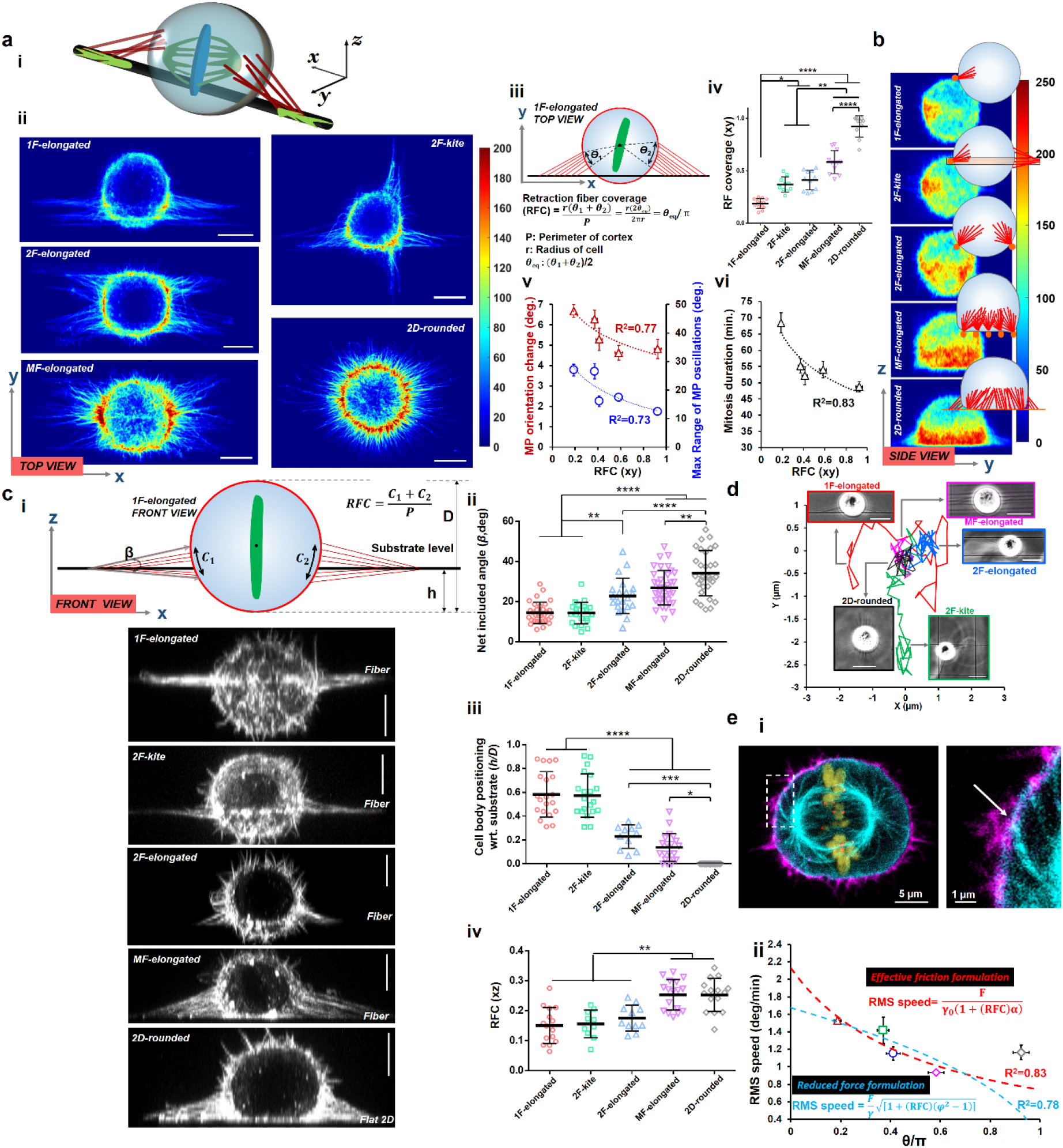
Retraction fiber-based stability and positioning of mitotic rounded cell bodies. (a,i) 3D schematic of rounded cell body held in place by retraction fibers in a single-fiber category, (ii) Average hotspots of retraction fibers from confocal microscopy of top view across various fiber configurations and 2D (n=10 per category), (iii) schematic describing retraction fiber coverage metric, (iv) retraction fiber coverage across fiber configurations and 2D in xy plane, (v) metaphase plate orientation and maximum range of oscillations decrease with increasing retraction fiber coverage, (vi) mitosis duration decreases with increasing fiber coverage. (b) Average hotspots of retraction fibers from confocal microscopy of side view across various fiber configurations and 2D (n=10 per category). (c, i) Representative images showing front view of mitotic cells across various fiber configurations and 2D, schematic shows the retraction fiber organization from the front view of cells, (ii) cell body positioning (h/D ratio defined in c(i)) across all fiber configurations and 2D, (iii) net included angle (β, defined in c(i)) across all fiber configurations and 2D, (iv) retraction fiber coverage in xz plane across all fiber configurations and 2D, (d) Representative cell body displacement profile during mitosis showing cells in 1F-elongated and 2F-kite shapes undergoing large displacements. (e, i) Representative image showing the mechanical linking of the astral microtubules and the cell cortex at the sites of the retraction fibers, Actin (magenta) microtubules (cyan), histone H2B (olive), and kinetochores Hec1(red). (ii) Model formulations based on increased effective friction and force for MP movements at higher retraction fiber coverage

Next, given our observations of different coverage of retraction fibers, we wanted to investigate the 3D shapes of mitotic cells, their placement with respect to the fiber axis, and their movement. Quantification of the mitotic cell heights and aspect ratios revealed that multifiber and flat 2D cases supporting the highest number of focal adhesion clusters and retraction fibers caused cell bodies to be non-spherical, i.e., the bodies had reduced heights and lower cell height-width ratios (**Fig. 2b, Fig. 1b**). Our observations are in excellent agreement with previous studies on flat 2D, which also report shorter cell heights than widths^47^. To relate the spatial arrangement of retraction fibers and cell body with respect to fiber axis (**Fig. 2c(i)**), we introduced new metrics: i) the number of retraction fibers associated with a single FA cluster, ii) location of rounded cell with respect to the substrate, and iii) net included angle (β), defined as the angular spread of the retraction fibers about their attachment sites. Quantification of the number of major retraction fibers emanating from individual FA clusters revealed an average of 10-12 retraction fibers per category (**Supplementary Fig. S3**). Interestingly, analysis of the cortical distribution of retraction fibers revealed that the retraction fibers were mainly positioned along the nanofiber plane in 1F-spindle and 2F-kite configurations. Cells in the 2F-elongated, MF-rectangular and flat 2D cases were positioned above the substrate, causing the retraction fibers to be organized at a steeper angle between the mid-cortical level of the rounded cell and the underlying substrate. A significant number of 2F-elongated cases were found to be entrapped between the two fibers at varying heights (confined cases excluded from this analysis and described in detail later in the manuscript). Thus, not surprisingly, the net included angle and the retraction fiber axial distribution was found to be significantly higher in MF-elongated and the 2D-rounded categories **(Fig 2c (ii))**.

Additionally, we observed that the rounded mitotic cell was positioned roughly in the mid-cortical level for 1F-spindle and 2F-kite shapes (h/D ratio~ 0.5-0.6). In contrast, cells on multiple fibers typically were positioned on top of the fiber plane (h/D ratio~0.1) similar to cells on Flat 2D (Fig. 2a(ii), **Fig. 2c(iii)**). Increased retraction fiber coverage directly affected the rounded cell body movement observed from high-speed timelapse videos (TOP view, **Supplementary movies 10-13**). On flat 2D and multifiber categories, the cell body was essentially locked in place (minimal movement), while 1F and 2F categories displayed significant cell body movement. Though unviewable from the FRONT view, we expect similar trends of cell body movement in 1F and 2F-kite due to h/D ratio of ~0.5 and reduced motion for 2F-elongated categories due to the mitotic bodies resting on top of the two fibers (low h/D ratio~0.22). Overall, we expect that 1F and 2F-kite mitotic rounded bodies undergo substantial 3D movement, including rotations about all axes.

Next, we developed a force/friction-based theoretical model to understand our experimental observations of metaphase plate stability. We present two formulations, with both models based on the force interactions between the actin cortex and astral microtubules (**Fig. 2e(i)**). The first formulation considers the higher effective friction experienced by the interacting microtubules in the cell cortex region covered by the retraction fibers. Thus the root-mean-square (RMS) angular velocity of the metaphase plate (quantified from distributions of the angular velocity, **Supplementary Fig. S4**) scales with the microtubule-cortex forces (F) as, 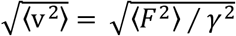, where *γ* is the effective friction coefficient, which is given by *γ*=*γ*0(1+RFC\**α*), where *α*>1 denotes the higher effective friction in the cortical region attached to the retraction fibers. Thus, the RMS speed of the metaphase plate depends directly on the retraction fiber coverage as 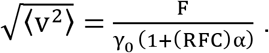.

The second formulation considers a reduction of the forces exerted by the astral microtubules on the cortex due to being “stuck” on adhesion complexes where the retraction fibers emanate. The central approximation in this formulation is that microtubules are less mobile when in contact with a cortex area with retraction fibers linked to it, by a factor 0 < φ < 1. Thus, the mean square force (microtubule-cortex) can be given as:

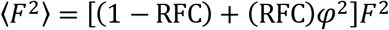

and the corresponding RMS speed 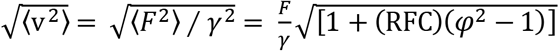

Fitting this expression to the experimental results demonstrates that φ is effectively zero, which gives the dependence of the MP RMS speed on the retraction fiber coverage in this limit: 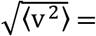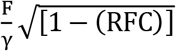.

Both of the proposed models fit our experimental data reasonably well (R^2^=0.83 and 0.78 respectively, **Fig. 2e (ii)**) and demonstrate how increasing the RFC can significantly reduce the overall movements of the metaphase plate.

Overall, we found that the spatial organization of retraction fibers in fibrous environments regulated the 3D positioning, shape, and movement of mitotic cells and led to varying levels of mechanical stability of the metaphase plate.

### Dynamics of daughter-cell positioning

Since 3D mitotic cell body positioning, metaphase spindle dynamics, and mitotic times governed the organization and density of retraction fibers, we inquired if the orientation of the cell division axis and the subsequent positioning of the dividing daughters were also affected. We quantified the orientation of the division axis with respect to the underlying fiber orientation. Interestingly, 1F-elongated cells, the least stable cells, demonstrated the widest spread (**Fig. 3a**) in their division axis orientation compared to the 2F-rectangular or MF-rectangular cell shapes. Consistent with average metaphase plate orientations, 2F-kite shaped cells showed a distinct peak (~ 67%) in the 30-60° region, indicating the division axis to be symmetrically located between the two orthogonal focal adhesion clusters. On Flat 2D substrates, due to the isotropic distribution of retraction fibers, the division axis distribution was primarily random, in agreement with the previous reports^26^.

**Figure 3:**
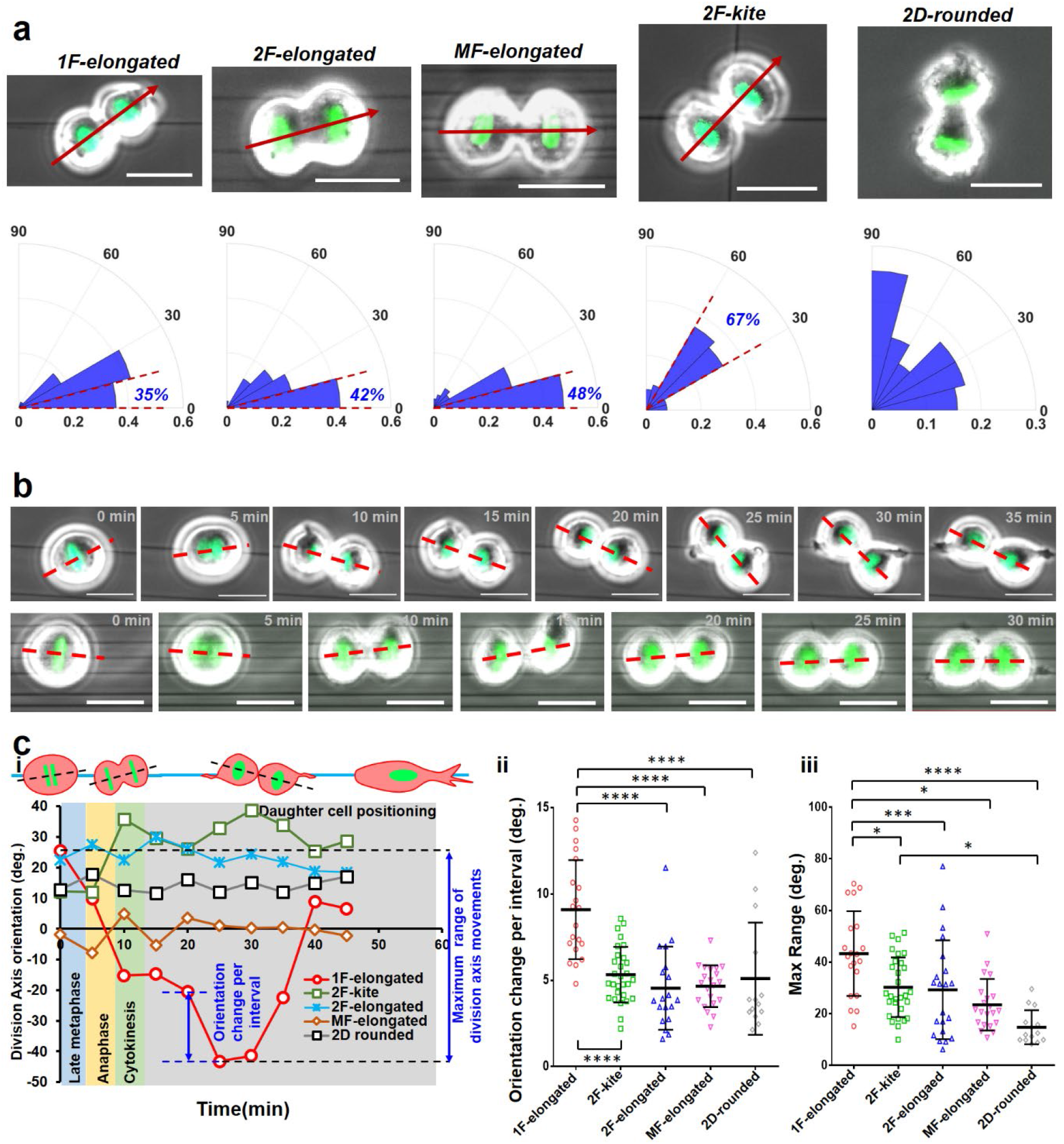
Division axis orientation and daughter cell positioning on suspended fibers: (a) Polar histograms of cell division axis orientations (marked by red arrow) in for different fiber geometries, absolute acute angle magnitudes of orientation are used, n=45, 41, 62,105 and 50 for 1F-elongated, 2F-elongated, MF-elongated, 2F-kite and 2D rounded, respectively. Percentage of cells in 0-15° sector for aligned geometries, and 30-60° for crosshatch geometry, (b) Timelapse images showing the representative dynamics of cell division axis (red dashed line) in single fibers versus multiple fibers. (c) i) Representative profiles showing the variations of division axis orientation and subsequent positioning of daughter cells, (ii,iii) Statistical comparison of the max range of division axis movements and orientation change per interval, for the different substrate categories.

Additionally, following mitotic exit by the formation of the cytokinetic furrow, we observed that the orientation of the cell division axis is highly dynamic and tracked it from late metaphase (see methods for defining cell division axis orientation) till the time taken by daughter cells to begin spreading (**Fig. 3b**). As with the previous analysis (Fig.1), we utilized two major parameters: the maximum range of orientation change in the total time period and average orientation change during individual imaging intervals (**Fig. 3c(i)**). In line with our analysis of the maximum 3D movement of mitotic 1F-elongated rounded bodies, we found that daughter cells of 1F-elongated cells were the most dynamic as they underwent the maximum degree of motion before spreading on fibers (**Fig 3c (ii, iii)**). High-speed timelapse microscopy demonstrated that the fluctuations in daughter cell positioning were primarily mediated by the retraction fibers (**Supplementary Movies 10-13**). Not surprisingly, mitotic rounded cells essentially locked in position through the multiple retraction fiber groups on multiple fibers (MF-rectangular shape) demonstrated significantly reduced fluctuations in daughter cell spreading. Overall, we found that retraction fiber-based stabilized mitotic cells and positioned the daughter cells following mitotic exit.

### Mitotic fidelity is influenced by fiber geometry

Our observations that controlling interphase cell shape and adhesion geometry can significantly affect the metaphase plate dynamics, mitotic duration, and daughter cell positioning led us to inquire if the mitotic errors were affected by fiber geometry as well. To this end, we synchronized divisions of HeLa cells by treatment and washout with the Cdk1 inhibitor RO-3306 and fixed mitotic rounded cells at metaphase^48,49^. We immunostained the rounded cells for different components of the mitotic spindles (**Fig. 4a**): microtubules (β-tubulin), kinetochores (Hec1), and the chromosomes (Histone H2B). We quantified the net length and width of the mitotic spindle (**Supplementary Fig. S5**). We did not observe differences between the overall spindle shape across fiber networks and conventional flat glass substrates. Since kinetochore stretching is directly associated with regulation of mitotic timing^50^, we wanted to estimate the extent of stretching by analyzing the inter-kinetochore separation distances(***δ*** in **Fig. 4b**). Our observations reveal that the inter-kinetochore separation distance was reduced in the 1F-elongated cells (**Fig. 4c**), thus, potentially explaining the slower mitotic progression in these cells (**Fig. 4d**).

**Figure 4:**
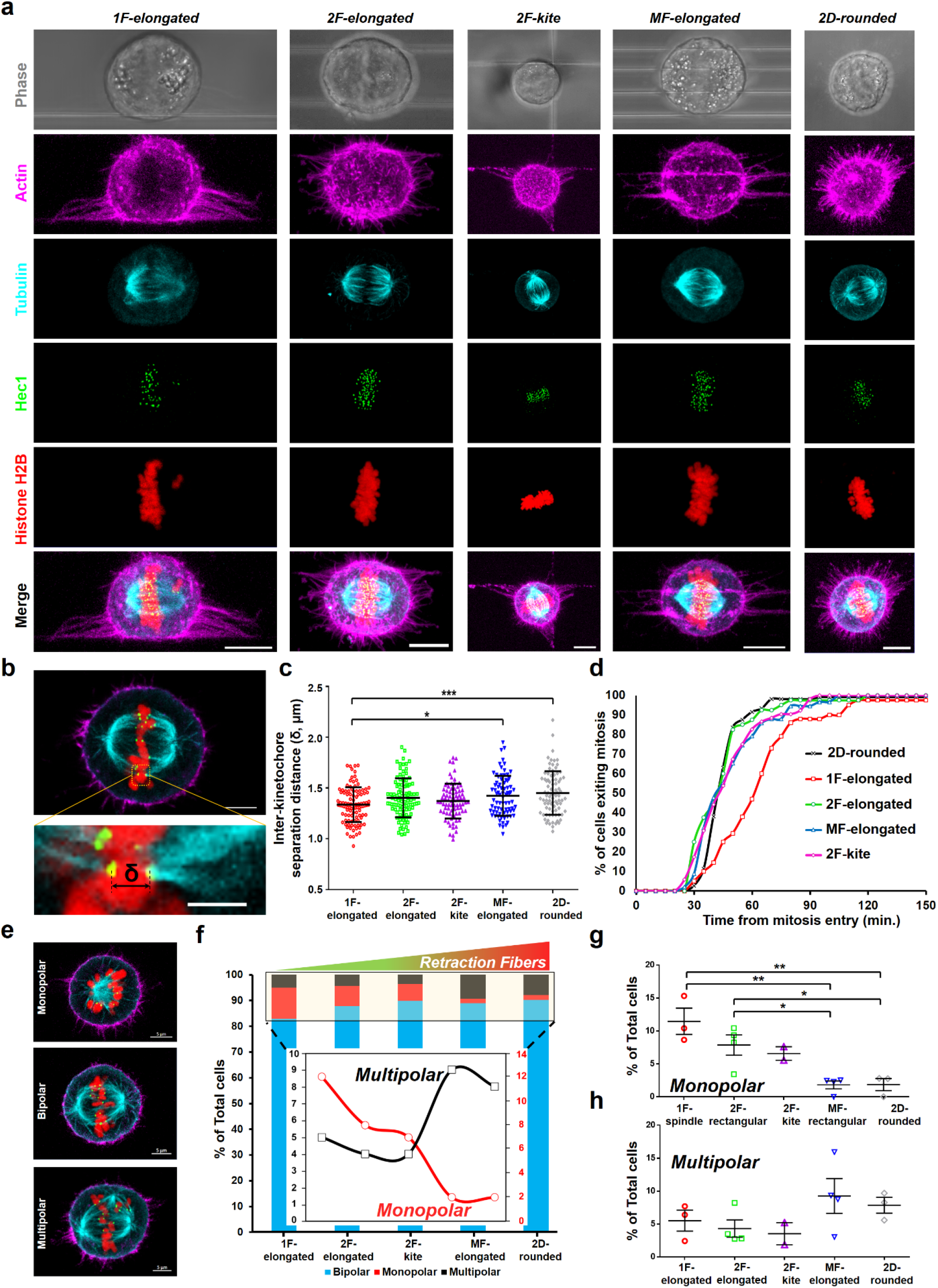
Mitotic spindles in different fiber geometries: (a) Cells are fixed at metaphase and stained for the actin cortex (magenta), microtubules (cyan), kinetochores (green) and chromosomes (red). Maximum intensity projected images are shown. Brightfield images show the respective fiber networks. (b) Representative image showing a pair of kinetochores attached to both ends of a single chromosome via microtubules. (c) Comparison of the inter-kinetochore separation distance between the different cell shapes. (d) Cumulative profiles showing the mitotic progression for the different categories. (e) Confocal images of the three major categories of mitotic spindles based on the number of observable poles. (f) Relative occurrence of monopolar, bipolar and multipolar spindles in the different cell shape categories. Inset shows the % of total cells in box area. Increased stability due to increased number of retraction fibers causes an increase in multipolar and a decrease in monopolar defects. (g,h) Comparison of monopolar and multipolar defects in the different substrate categories.

Next, we wanted to investigate if the mitotic spindle assembly was affected by the underlying fiber geometry. Similar to the previously reported classification of mitotic spindles, we also observed three major types of outcomes (**Fig. 4e**): i) bipolar (2 symmetrically positioned spindle poles for normal chromosome segregation), ii) monopolar (1 spindle pole, defective chromosome segregation), and iii) multipolar (at least three spindle poles, defective chromosome segregation)^51^. To understand the relative occurrence of these spindle types, we sampled at least 300 cells (from multiple independent experiments) for each cell shape category. We observed significant differences in the relative occurrences of the mitotic spindles. Increasing the stability during mitosis (increasing the number of retraction fibers) resulted in an increased percentage of bipolar segregation (**Fig. 4f**). Contrarily, decreasing the stability during mitosis (1F-elongated and 2F-kite) resulted in a significantly higher percentage of monopolar spindles (**Fig. 4f, 4g**). Unexpectedly, we found increased incidences of multipolar defects at high stability during mitosis (MF-elongated and 2D-rounded, **Fig. 4h**). Overall, we found that retraction fiber-mediated stability contributed to mitotic spindle integrity and chromosome segregation outcomes.

### Confinement enables measurement of mitotic forces and causes tilt of mitotic spindles

Mitosis proceeds with cells rounding up and exerting outward forces, and natural ECM environments impose confinement. We wanted to inquire about the role of confinement in mitotic progression. We found that the 2F-elongated cells often got trapped (confined) within the two parallel fibers resulting in an outward push of the fibers (**Fig. 5a, Supplementary movies 14,15**). To quantify the extent of confinement, we analyzed the positioning of the rounded cell with respect to the fiber plane from cross-sectional side views (yz, **Fig. 5b(i)**). We found that suspended fibers were located roughly at the mid-cortical level (h ~ H/2) in a significant proportion of cells (~35%). We defined confined cases (h/H~0.4-0.6), which from timelapse microscopy corresponded to maximal outward deflection of fibers of at least 2 μm. We categorized cells as weakly-confined for h/H values ranging from 0.2-0.4 and 0.6-0.8 and unconfined (nearly sitting on top of the fibers) for lower (<0.2) or higher (>0.8) h/H values. Interestingly, we found that mitotic cells under confinement were significantly taller (increased cell height, **Fig. 5b(ii)**) with a higher aspect ratio (**Fig. 5b(iii)**) as compared to their weakly-confined and unconfined counterparts.

**Figure 5:**
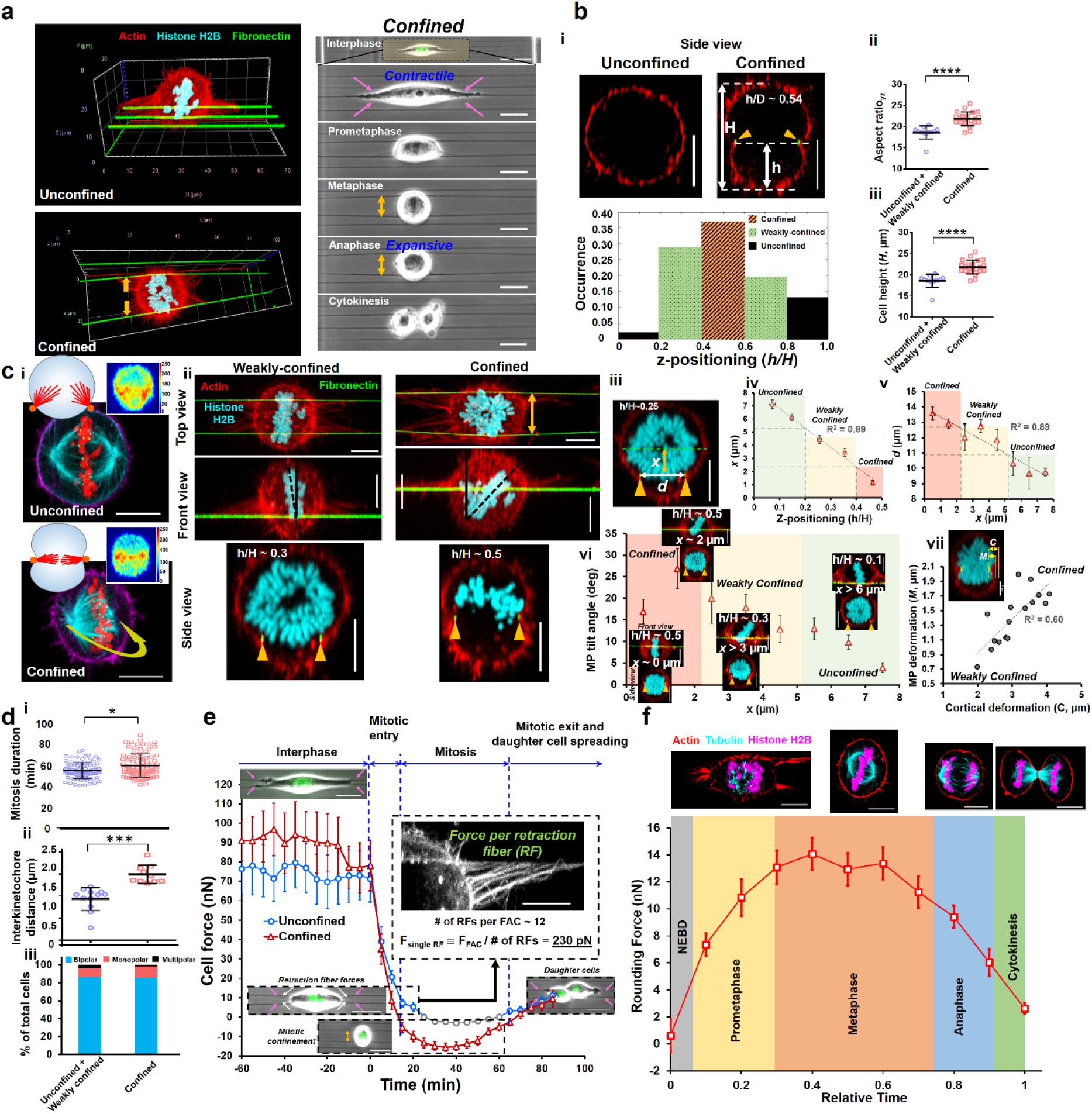
Cell division under confinement. (a) Isometric views of a representative cells positioned on top of the fiber network (unconfined category) and confined between 2-fibers (confined category). Actin (red), fibers (green), chromosomes (cyan). During mitosis, the contractile forces change direction from inward (pink arrows) to outward directions as cell balls up and pushes the fiber outwards (yellow arrows). Scale bars 20 μm. (b, i) Side view (yz) of a representative confined cell. Actin (red) and fibers (green) are indicated by yellow arrowheads. Z-positioning of the cell with respect to the fiber plane is defined in three main categories according to ratio of h/H, where *h*: is height of rounded cell below the fiber plane, and *H* is the net cell height. *h/H* values of 0.4-0.6 are considered confined, 0.2-0.4 and 0.6-0.8 are considered weakly confined, and 0-.2 and 0.8-1.0 as unconfined. Scale bar is 10 μm, (ii) Comparison of aspect ratio and cell height between the unconfined+weakly confined and confined categories. (c, i) Confinement causes rotation of the mitotic spindles inside the cell, while the retraction fiber coverage (inset) remains similar, (ii) Rotation of the mitotic spindles causes a tilt of the metaphase plate (seen from confocal front views), (iii) Positioning of the metaphase plate with reference from the positioning of fibers on cell cortex is defined by vertical distance (*x*) from confocal side views. The distance between the two confining fibers is denoted by *d*. (iv, v) cell z-positioning (*h/*H) against *x* shows a linear trend to define three categories of confinement in terms of distance *x*: confined ~ 0-2.2 μm, weakly-confined ~2.2-5.5 μm, and unconfined ~> 5.5 μm. (v) Plotting the interfiber spacing distance ‘*d*’ vs. *x* shows a linear trend-increasing *x* denoting unconfined categories leads to lower *d*, (vi) Plotting metaphase tilt angle vs. *x* shows that in confined categories, at low values of *x*, fiber pinching can cause a low tilt angle, while slight increase in *x* values causes a significant increase in tilt angle. As *x* increases, the tilt angle decreases. In unconfined categories, the metaphase tilt angle approaches zero angle, (vii) Metaphase plate deformation (*M*) increases with cortical deformation (*C*). Measurement methods are shown in the inset image. (d) Effect of confinement causes a drop in mitotic times (i), increase in interkinetochore distance (ii), and a decrease in multipolar defects. (e) Force profiles of confined and unconfined cells. Inset shows analysis of approximating tension in each retraction fiber (n= 20 per category). Forces for the retraction fiber analysis are taken from the unconfined force profiles and divided by four to obtain force per focal adhesion cluster length. (f) Force profiles during cell rounding and mitosis are normalized for time taken from NEBD to cytokinesis (n=23) with representative top views of cells undergoing division.

Since confinement caused mitotic cells to be of a high aspect ratio, next, we interrogated if it affected the organization of mitotic spindles. We found that confinement caused the mitotic spindles to rotate inside the cell body (**Fig. 5c(i)**), matching tilted spindles found during division in epithelial tissues^40,52^. Next, we sought to quantitate the tilt of the metaphase plate in confined conditions. Averaging the intensity plots from confocal images showed retraction fiber hot spots localized at the equatorial plane, similar to unconfined cases, thus suggesting a reduced role of retraction fibers in causing the rotation. Analyzing the three views through confocal microscopy revealed the metaphase plate to be significantly tilted in confined cells (**Fig. 5c(ii)**). To understand this further, we looked at the contribution of pinching of fibers from the equatorial plane (distance *x*) and the distance between the external fibers (*d*, **Fig. 5c(iii)**). Unconfined cells sitting on top of external fibers would have the highest *x* value (**Fig. 5c(iv)**), coinciding with the lowest *d* values (**Fig. 5c(v)**). Our analysis showed that confined cells had *x* ranging from ~0-2, weakly-unconfined from ~2-5, and higher values signifying unconfined cases. In doing so, we quantitated the MP tilt angle with increasing values of *x* (**Fig. 5c(vi)**). Unconfined cases of high *x* values were, as expected, found to have the lowest MP tilt. In confined cases, the cell body became two distinct symmetric lobes with the confinement causing a pinch in the metaphase plate (**Fig. 5c(vii)** and **Supplementary figure S6a**).

We found the MP tilt to be sensitive to the location of the pinch by the external fibers. At values of *x* close to zero, we found the MP tilt angle to be close to zero and low tilt values. However, as *x* increased, we found a significant increase in MP tilt values, thus indicating the system's sensitivity to the location of confinement from external fibers. At *x* values approaching ~2, even though the cell body was divided into two distinct lobes, the metaphase plate was pushed preferentially in one lobe, thus causing the higher tilt. With further increase in *x* signifying weakly-unconfined categories, the cell body became asymmetrically lobular. The metaphase plate was found to be contained in the larger lobe and had a reduced tilt angle. We measured the deformation of cell cortex and MP and found that increasing cortical deformations corresponded with an increase in the MP pinch (Fig. 5c(vii)). These findings suggest that a combination of intracellular pressure and convex geometry of cortical deformation play a role in controlling the MP tilt angle. Furthermore, if MP were a perfectly rigid disc that can rotate to avoid bending of the confining fibers, then the tilt would be predicted to be larger than the observed (**Supplementary figure S6b**), supporting our observations that MP is deformable (Supplementary figure S6a). Incidentally, confinement caused a significant drop in metaphase plate (MP) fluctuations (**Supplementary figure S6c**), which coincided with faster mitotic progression (**Fig. 5d(i)**)-findings that are in line with increased stability (Fig. 1), and suggesting that the stabilizing external fibers overrode retraction fiber cues in confined cells. We also found an increase in the angle of the division axis (**Supplementary figure S6d, S6e, Supplementary Movie 16**) which can be probably attributed to the tilted orientation of the mitotic spindle. The increase in angle caused the daughter cells to spread on adjacent fibers (Supplementary Movie 16). Confinement also caused an increase in interkinetochore distance (**Fig. 5d(ii)**). In contrast to retraction fiber-based stability mitotic errors (Fig. 4f), we found a slight increase in monopolar defects in cells with rotated mitotic spindles (**Fig. 5d(iii)**).

We took advantage of entrapment of the mitotic cell between 2-parallel fibers (2F-rectangular shape during interphase, **Fig. 5e**) to estimate the rounding forces during mitosis using a single point force model in Nanonet Force Microscopy^42,43,53,54^ (NFM, appendix). During interphase, cells are in their natural contractile state and thus deflect the fibers inward (pink arrows). However, following mitotic entry in confined conditions, as cells progressively round up, they push out the fibers to cause distinct outward deflections (Fig. 5e, yellow arrows). In the case of weakly-confined and unconfined cells, we observed the balled-up cells held in position by retraction fibers (white arrows) while still deflecting the external fibers. Knowing the average number of retraction fibers (average ~11 per FAC and Supplementary Figure 3) and the forces exerted allowed us to approximate the tension in each retraction fiber (~230 pN) by assuming retraction fibers to be springs in parallel. Consistent with previously reported literature^47,55^, we observed a steady increase in the rounding forces from nuclear envelope breakdown, with peak mitotic forces of ~12-14 nN averaged over multiple cells. Following mitotic exit (anaphase completion), rounding forces dropped sharply, with outward fiber deflections becoming negligible. Finally, with daughter cell formation and spreading, fiber deflections were again inward, indicating transition into the contractile state.

Overall, external mechanical confinement by fibers rotated the mitotic spindles, caused significant tilt of metaphase plate, and accelerated mitotic progression.

## Discussions

Here, we inquired about the role of mechanical stability in establishing mitotic rules for cells attached to suspended and physiologically relevant ECM-mimicking nanofibers of well-controlled geometries. We designed aligned and suspended fiber networks with tunable inter-fiber spacing and organization to understand the interplay of interphase cell shape and adhesion organization. Spacing fibers apart by more than 20 μm led to cells attached to single fibers in elongated spindle shapes (1F-elongated) at two major focal adhesion clustering (FAC) sites at the cell peripheries. 2F-elongated category represented fibers at a separation distance of 10 μm resulting in cells attached to both fibers through four periphery FACs. In the MF-elongated category, fibers were spaced apart by ~5 μm resulting in cells attaching to three or more fibers (six peripheral FACs or more). We achieved precise control of cell elongation on aligned fibers, with decreasing aspect ratio as the cells were connected to more fibers (1F>2F>MF). Crosshatched fibers constrained cells to kite shapes (2F-kite) with an aspect ratio similar to flat 2D control.

Proper cell rounding during mitosis plays a critical role in chromosome capture, spindle stability, and allowing the spindle to locate the center of the cell accurately^56^. Rounded mitotic cells retain their ‘memory’ of interphase shape through force-bearing retraction fibers^10^ or *mitotic* protrusions^13^. The strategy to precisely control fiber networks allowed us to investigate mitotic progression in rounded cell bodies due to stability provided by the actin-rich retraction fibers. We found that the organization of retraction fibers resulted in the shaping and positioning of mitotic rounded cell bodies. We grouped cells by their aspect ratios from the side view of confocal images: high aspect ratio (1F, 2F-kite, and 2F) and low aspect ratio (MF and flat 2D). In the high aspect ratio category, cells achieved near-perfect circular shapes and were positioned on either side of the fiber in the Z direction. However, basal surfaces (flat 2D glass and fibers) flattened the base of rounded cells in the low aspect ratio categories, resulting in cells positioned above the basal plane. Our findings agree with previous studies demonstrating retraction-fiber-driven sagging of rounded mitotic cells on flat surfaces, resulting in a flattened morphology^47,57^. Confinement in 2F-elongated cells had the remarkable influence in shaping cells with distinct two lobes as cells pinched the cortex at the equatorial plane and a single lobe of asymmetrical shape positioned above the fiber axis due to pinching happening away from the equatorial plane. The differences in mitotic cell shapes and their positioning can potentially explain why multipolar defects are more prevalent in the slightly flattened mitotic cells on multiple fibers or flat surfaces, while monopolar defects are more common for the highly circular mitotic cells^56^.

Consistent with observations on elongated cells on micropatterns, cells on aligned fibers divided along the fiber axis^3^. Our observations that the metaphase plate can undergo significant levels of oscillations are in agreement with previous studies, which also report similar movements throughout the division process^45^. An increase in retraction fibers (retraction fiber coverage) resulted in lower MP oscillations and faster mitotic times, in agreement with a previous study demonstrating that a loss of retraction fibers mediated by knockdown (KD) of β_5_-integrins led to random orientations of the mitotic spindle and delayed mitosis^15^. Here we show that the amount of cortical coverage by retraction fibers is determined by the geometry of the cell-ECM adhesions, which thereby critically affects the fidelity of cell division.

Recent studies have demonstrated how astral microtubules existing at either end of the mitotic spindle are mechanically linked to the cell cortex and the retraction fibers via the integrin-dependent cortical mechanosensory complex^6^. The tension in the astral microtubules depends upon the contact angle between the cell surface and microtubules. For spherical cells, microtubules contacting the cortex at the shallowest angle exert the highest force^58^, suggesting the possible role of friction in establishing the forces that control mitotic spindles. Our theoretical formulation of the mitotic spindle stability is based on mechanical interactions/links between the astral microtubules and the cortical complexes connected to retraction fibers (integrins and other key players of the cortical mechanosensory complex, including caveolin-1, FAK, LGN-NuMA, dynein, and MISP, to the cell cortex^6,7,59^), which presumably contribute to increased friction. Our model, therefore, predicts a constraining effect on metaphase plate movements at higher cortical coverage by retraction fibers.

Mitotic cell rounding is associated with a multi-fold increase in the cortical tension^60^ and the intracellular hydrostatic pressure^47^, manifesting outward expansive forces required to undergo cell rounding within confined microenvironments. Failure to counteract mechanical confinement can lead to spindle defects^61^, delayed cell division^62^, and even cell death. Several sophisticated methods have been developed to study mitosis, and below, we highlight key differences observed in suspended fibrous environments:

First, we find differences in daughter cell placement compared to flat 2D. It has been shown that for cells attached to flat 2D substrates, daughter cells spread precisely according to the mother cell adhesive patterns. Thus, the most elongated cells will have the slightest deviation in daughter cell spreading directions^5^. In contrast, we find the most elongated shape (1F-elongated with interphase aspect ratio~15) and kite-shapes on crosshatches (2F-kite with interphase aspect ratio ~1) had the widest spread in the cell division axis orientation. The cells on single fibers (1F-elongated) have two sets of retraction fibers originating at the two peripheral FACs. In the 2F-kite category, the interphase cell is attached to four fibers (four FACs), but during mitosis, the rounded cell is located across two crossing fibers (two orthogonally located FACs). In both 1F-elongated and 2F-kite categories, the two sets of retraction fibers cause the rounded cell bodies to have significant movement in 3D during the mitosis process resulting in increased metaphase plate movements and mitosis duration times. Cells on multiple fibers (MF-elongated) match the outcomes from 2D flat control. Thus, we conclude that in fibrous environments, the interphase cell elongation factor is a poor predictor of daughter cell positioning.

Second, we find differences in the magnitude of forces measured during mitosis. Cell rounding forces during mitosis of HeLa cells measured using AFM cantilevers have shown peak forces at the metaphase to be in the range of ~40-90 nN depending upon the height of rounded cell^63–65^. The cell body, in these measurements, is constrained between two flat planes (AFM cantilever and flat basal substrate). In another study, outward mitotic forces were measured using two independent methods (AFM and vertical cantilevers) for MDCK line^55^. The authors found that force measurements using vertical cantilevers were significantly lower than those obtained by AFM methods (3-17 nN from vertical cantilevers vs. 43 nN using tipless AFM cantilever). The authors also tested HeLa cells and found maximum outward mitotic forces using vertical cantilevers to be ~15 nN. Outward rounding forces measured using our method (~12-14 nN) match those obtained using the vertical cantilever system. Furthermore, our rough measurements of retraction fiber tension (~230 pN) using springs in the parallel analogy are in excellent agreement with previously reported values using optical tweezers (250 pN)^10^. Extending these measurements provides us a rough estimate of ~ 2.5 nN tension per focal adhesion cluster, which translates to a retraction fiber-based tension of approximate 5 nN (1F-elongated), 10 nN (2F-elongated), 5 nN (2F-kite), and greater than 15 nN for cells on three and multifiber systems. Thus, 1F-elongated and 2F-kite are held the weakest in position by retraction fibers resulting in maximum cell body movement during mitosis. In our method, the cells before anaphase have near-perfect circularities, thus suggesting an unconfined mitotic process. In this study, our force measurement system (nanonet force microscopy, NFM) uses suspended 250 nm diameter fibers at least 350 μm in length and fused at the base of larger 2000 nm diameter fibers. In such a configuration, the fibers can be visualized as guitar strings capable of undergoing large deflections. Cell-fiber forces increase with the diameter of fibers^42^. Providing a mix of diameters (small and large) in the 2F category can be a powerful tool to explore the role of anisotropic confinement forces in mitosis in future studies^42,66^.

Third, we find differences in the role of confinement in mitotic outcomes. In the vertical cantilever force measurement system, the authors show that cells that cannot generate sufficient pressure to overcome confinement or cannot escape confinement have tilted mitotic spindles ^55^. Another study in 3D gels has demonstrated how forces can shape the mitotic spindle through buckling of microtubules^9^. In the case of cells confined (entrapped) between fibers (2F-elongated), the rounding of cell bodies during mitosis occurs in a constrained manner but with near-perfect spherical shapes due to the ability of fibers to undergo large deflections. However, we find that confinement occurring at the equatorial planes causes the mitotic spindles to be tilted is slightly greater than the tilt values occurring naturally in epithelia (~6 degrees from the epithelial plane)^4,40^. Confinement occurring a few micrometers away from the equatorial plane is associated with the highest tilt, matching values reported with loss of cadherins^40^. Rounded cells that are pinched away from the equatorial plane, a ratio of h/H > 2, signifying weakly confined and unconfined states, have lower mitotic tilt angles and can be considered to resemble epithelial cells escaping confinement through interkinetic nucleus migration (INM)^67,68^. While presently we do not have a direct method of controlling the level of entrapment, we envision that volumetric imaging of entrapped cells coupled with advancements in fiber spinning technologies capable of depositing adjacent 2F fibers with a height offset (1-5 μm) can be valuable to control spindle angles to study confinement-based outcomes (stratification, sprouting, maintenance, growth, and tumoricity^41^).

In conclusion, we demonstrate the utility of our suspended fiber-based method to develop new knowledge regarding mitotic progression in cells of different shapes. To a high degree, our method captures mitotic outcomes presumably occurring in cells sparsely dispersed in loose interstitial ECM. Additionally, under certain conditions of confinement, it replicates cell division outcomes in tissues. We show the stability of the rounded cell body as a critical metric in mitotic outcomes. The dramatic switch between monopolar and multipolar defects by simply changing the number of fibers highlights the importance of various arrangements of the ECM environment. Future investigations of ECM-like fiber networks with different properties (fiber diameter, stiffness, architecture, and adhesivity) should allow us to gain a deeper understanding of the ECM-mediated regulation of cell division, which will likely impact pathophysiological outcomes.

## Materials and Methods

### Manufacturing nanofiber networks

Suspended fiber networks were generated from solutions of polystyrene (MW: 2,000,000 g/mol; Category No. 829; Scientific Polymer Products, Ontario, NY, USA) dissolved in xylene (X5-500; Thermo Fisher Scientific, Waltham, MA, USA) at 7 wt%, using our previously reported non-electrospinning Spinneret based Tunable Engineered Parameter (STEP) technique^38,39^. Horizontal arrays of 250 nm fibers with inter-fiber spacings between 3-25 μm were deposited on large diameter (~2 μm) vertical support fibers placed ~ 350 μm apart. Support fibers were generated from 5 wt% solutions of polystyrene (MW: 15,000,000 g/mol, Agilent Technologies) dissolved in xylene. Crosshatch networks were prepared with orthogonal layers of 250 nm fibers with an inter-fiber spacing of ~50 μm. Fiber networks were bonded at intersection points using a custom fusing chamber.

### Cell culturing and cell division synchronization

HeLa cells expressing Histone H2B GFP were cells were cultured in Dulbecco’s modified Eagle’s medium (Invitrogen, Carlsbad, CA) supplemented with 10% fetal bovine serum (Gibco, Thermo Fischer Scientific) in T25 flasks (Corning, Corning, NY, USA) and maintained at 37°C and 5% CO_2_ in a humidified incubator. Nanofiber networks were first sterilized with 70% ethanol for 10 minutes, followed by functionalization with 4 μg/mL of fibronectin in PBS (Invitrogen, Carlsbad, CA). For select imaging experiments, fibers were coated with rhodamine-conjugated fibronectin (Cytoskeleton Inc.).

To synchronize divisions, cells were treated with 9 μM of the Cdk1 inhibitor RO-3306 for 20 h. Cells were subsequently released for division after 2 times wash with complete culture media. To capture cells at metaphase, paraformaldehyde (4%) fixation was performed ~ 1 h following drug washout.

### Live imaging

Time-lapse optical imaging was performed every 5 minutes for extended periods of time (~12-16h) to track multiple cell division events. Imaging was performed with a 20x 0.8 NA objective in a Zeiss AxioObxerver Z1 microscope. GFP fluorescence was captured using a FITC filter set. Experiments were performed under incubation conditions of 37°C and 5% CO_2_ (Zeiss, Oberkochen, Germany). To capture cell body movement during mitosis, imaging was conducted at 1-5 s interval at 40x 0.75 NA objective.

### Immunofluorescent staining and imaging

Cells were fixed with 4% paraformaldehyde for 15 minutes. Following 2 times PBS wash, permeabilization was performed with a 0.1% Triton X-100 solution. Permeabilized cells were washed with PBS (2x) and blocked with 5% goat serum (Invitrogen, Grand Island, NY) for 45 minutes. Primary antibodies were diluted in an antibody dilution buffer consisting of PBS with 1% Bovine Serum Albumin and Triton-X 100, and stored overnight at 4° C. Primary antibodies include Anti-beta tubulin (1:500, mouse monoclonal, 2 28 33, Invitrogen), Anti-Hec1 (1:1000, human monoclonal) and Anti-phospho-Paxillin (1:100, rabbit polyclonal, pTyr31, Invitrogen). Secondary antibodies diluted in antibody dilution buffer was added along with the conjugated Phalloidin-TRITC (Santa Cruz Biotechnology) or Alexa Fluor 647 Phalloidin (1:40-1:80, Invitrogen), and stored in a dark place for 45 minutes. Secondary antibodies include donkey anti-human IgG Alexa Fluor 555 (1:600), Goat anti-mouse IgG Alexa Fluor 405 (1:500, Invitrogen) and Goat anti-mouse IgG Alexa Fluor 647 secondary antibody (1:500, Invitrogen). Confocal microscopy was performed using a laser scanning confocal microscope (LSM 880, Carl Zeiss Inc.) with optimal imaging settings and z-slice thicknesses ranging from 0.36-0.5 μm.

### Cell shape metrics

To quantify cell shape during interphase, manual outlining of phalloidin stained cells were performed in Image J (NIH; https://imagej.nih.gov/ij/) and aspect ratio (cell length by width) was computed using the Bounding Rectangle function. Cell circularity is defined by 4πA/P^2^, where A is spread area (μm^2^) and P is perimeter (μm) of the cell. A circularity value of 1.0 was achieved for perfectly circular cells. For elongated or contoured shapes, this value is reduced and falls between 0 and 1.

To compute cell heights during mitosis, complete z-stacks were processed in the Zen Blue (Carl Zeiss Inc.) and orthogonal cross-section views (xz and yz) generated using the ‘Ortho’ function were used.

### Quantification of cell division axis orientation

Orientation of cell division axis (corresponding to Fig. 3a analysis) is quantified at the late anaphase stage. The direction of the axis is taken along the line joining the centers of the two cell lobes corresponding to daughter cells. Once daughter cells have formed, the division axis (Fig. 3 b,c) is defined by the line joining the centroids of the two daughter cells. The division axis orientation is taken with respect to the undeflected horizontal fibers at all times.

### Scoring of mitotic defects

Cells stained for microtubules and DNA were visually inspected under a 40x objective to categorize as bipolar, monopolar or multipolar based on the number of spindle poles. At least 300 cells over 3-4 independent experiments were considered for each fiber category and Flat glass and the average proportion of observed bipolar, monopolar and multipolar cells are reported.

### Quantifying cell rounding forces

Outward deflections of nanofibers during cell rounding were converted to force (nN) values using our previously reported Nanonet Force Microscopy^42^. Briefly, fibers are modelled as Euler Bernoulli beams with fixed boundary conditions, and subjected to point loads at cell-fiber contact regions (see appendix).

### Statistical analysis

Statistical analysis was performed in GraphPad Prism (GraphPad Software, La Jolla, CA, USA) software. Statistical comparison among multiple groups were performed using one-way ANOVA along with Tukey’s honestly significant difference test. Pairwise statistical comparisons were performed using Student’s t-test. Error bars in scatter data plots indicate standard deviation. *,**,***,**** represent p< 0.05, 0.01, 0.001 and 0.0001 respectively.

## Author Contributions

ASN conceived and supervised the research. ASN, AJ, JD, and NG designed research. JD generated HeLa cells lines with tagged Histone2B. AJ, DZ, and JW conducted experiments. RK, AJ, and ASN developed the nanonet force microscopy force formulation. NG developed and implemented the theoretical model of retraction fiber stability. AJ, DZ, JW, ASN, JD, and NG analyzed data. AJ wrote the manuscript. All authors contributed to the editing of the manuscript.

## Acknowledgments

ASN acknowledges partial funding support from National Science Foundation (NSF, Grant No. 1762468). ASN acknowledges the Institute of Critical Technologies and Science (ICTAS) and Macromolecules Innovative Institute (MII) at Virginia Tech for their support in conducting this study. N.S.G. is the incumbent of Lee and William Abramowitz Professorial Chair of Biophysics, and this research was supported by the Israel Science Foundation (Grant No. 1459/17). JGD acknowledges funding support from the NIH (R35GM130365).

