## Supplementary Information for "Mitotic Outcomes in Fibrous Environments"

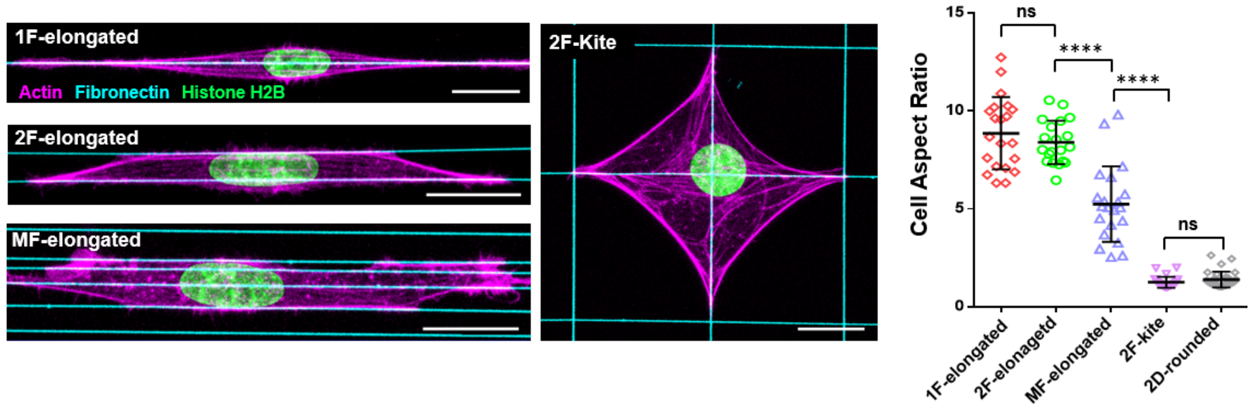


**Supplementary Figure S1: Cell shapes during Interphase:** Representative stained images and cell aspect ratio quantification corresponding to 1F-elongated, 2F-elongated, MF-elongated, 2F-kite and 2D-rounded

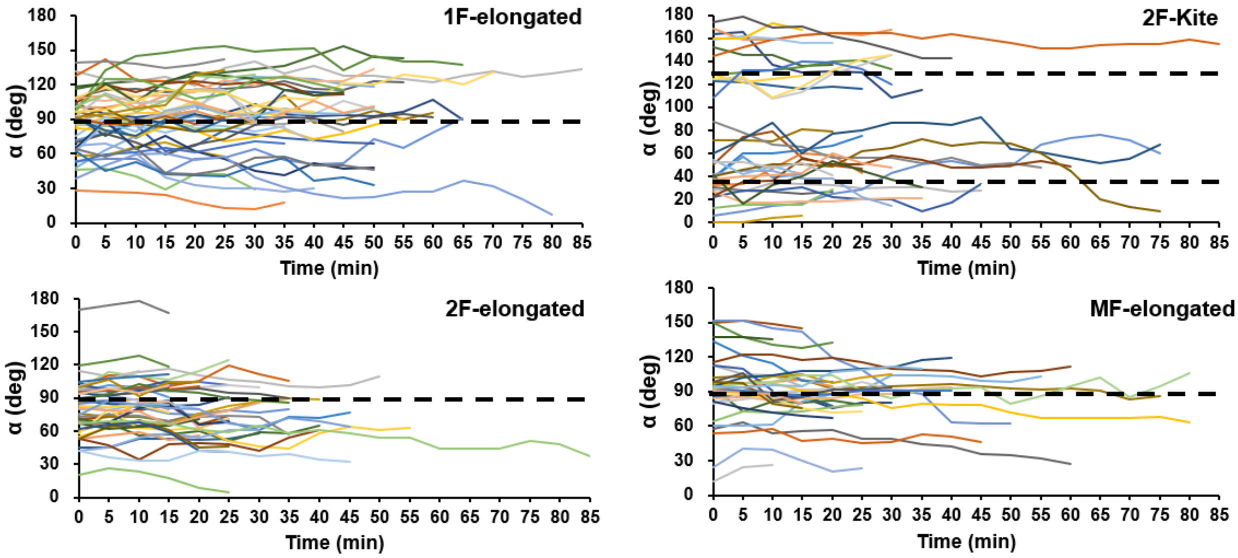


**Supplementary Figure S2: Compilation of individual profiles showing temporal dynamics of metaphase plate movements for the different fiber geometries**

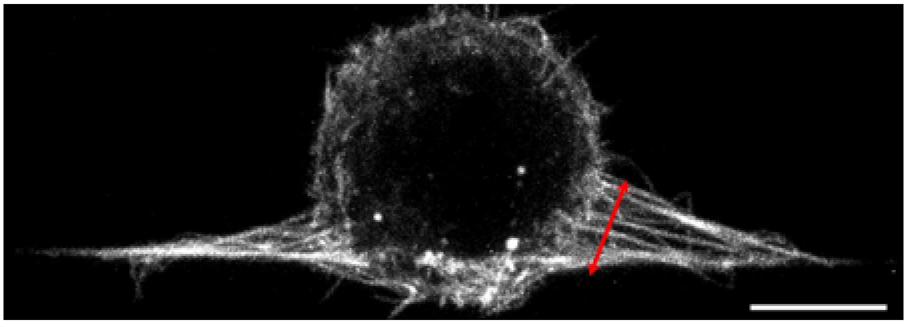

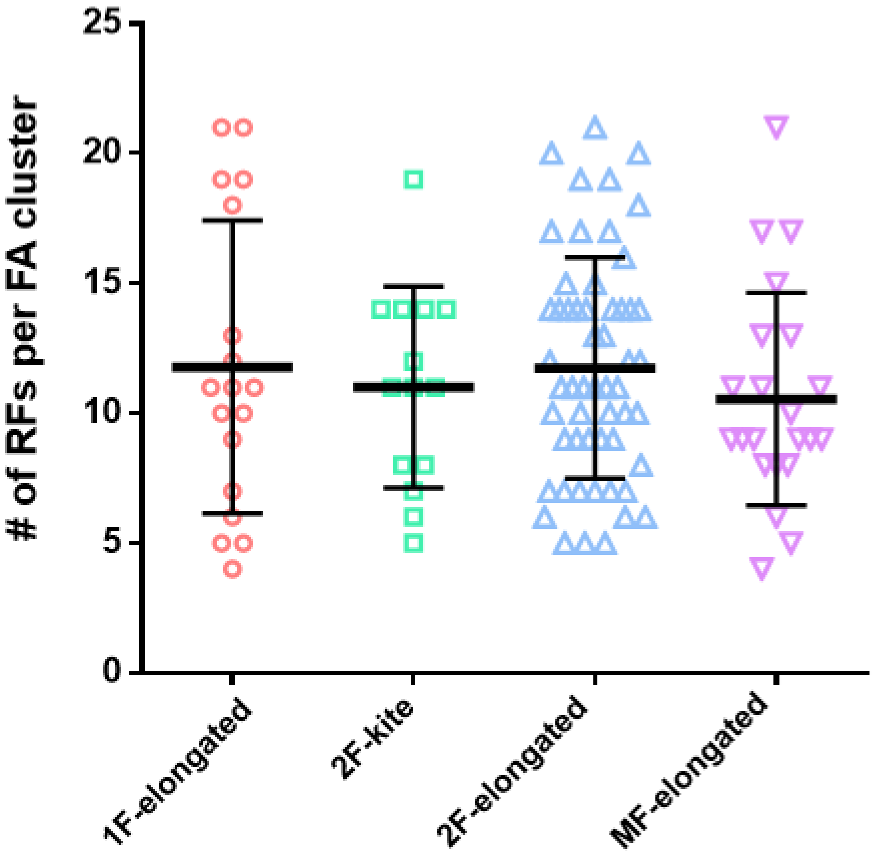


**Supplementary Figure S3: Quantification of number of retraction fibers associated with each focal adhesion cluster for the different cell shapes.** Red arrow in representative actin-stained image demonstrates the retraction fibers

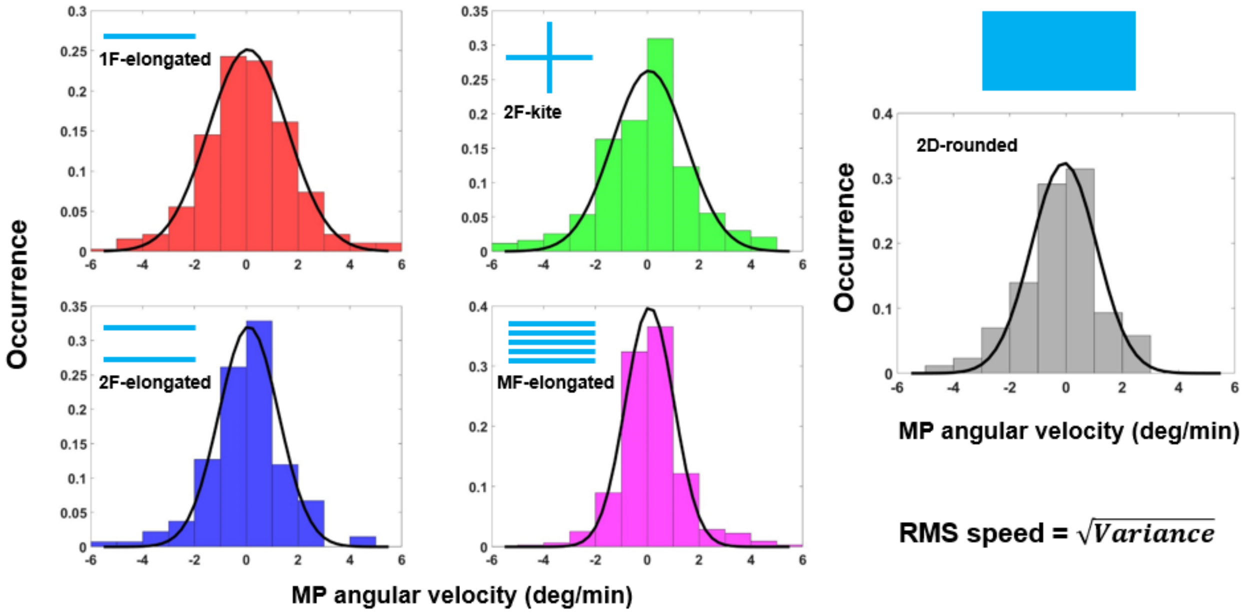


**Supplementary Figure S4: Probability distributions of the metaphase plate angular velocity for the different substrate categories**

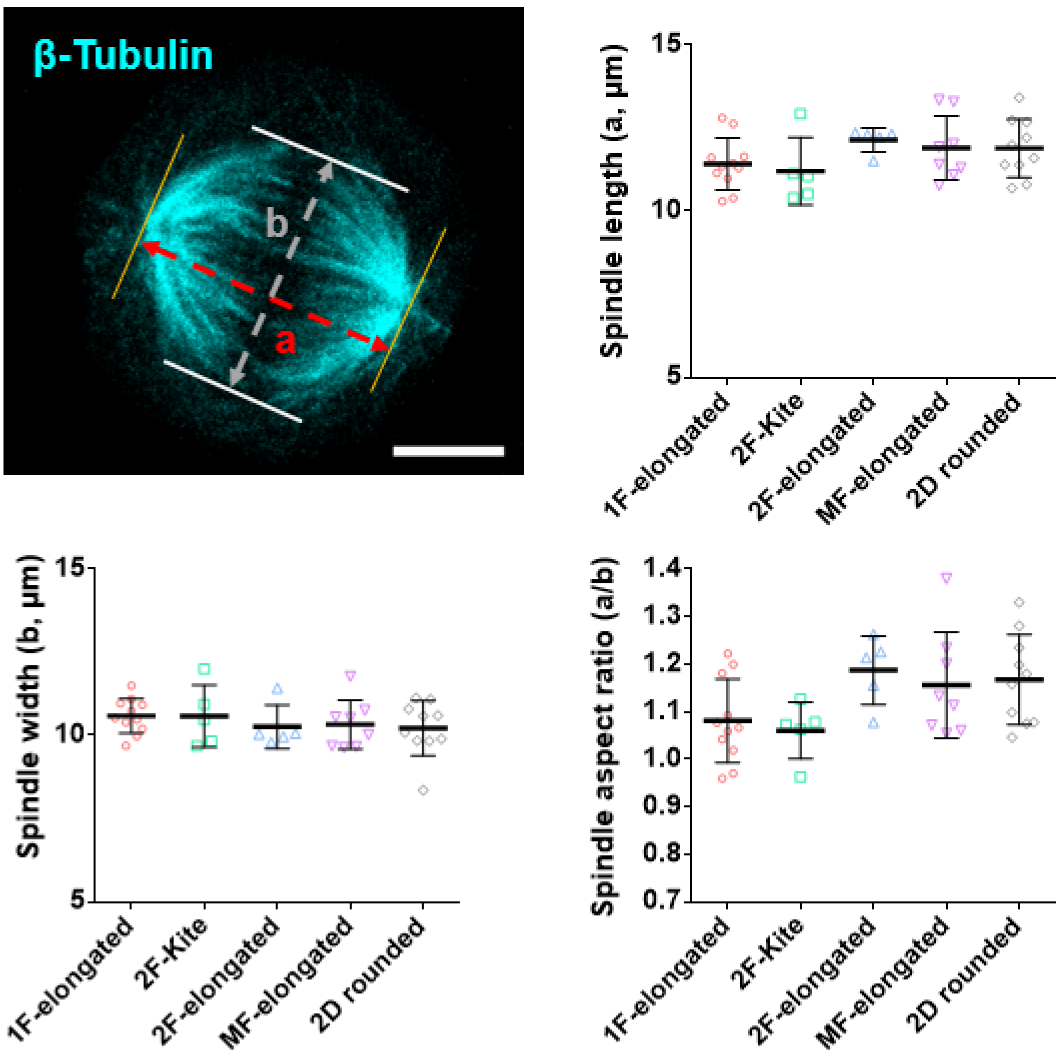


**Supplementary Figure S5: Comparison of mitotic spindle shape over different substrate categories.** The length and width of the mitotic spindle is shown by a and b in the representative stained image. Aspect ratio is defined as a/b.

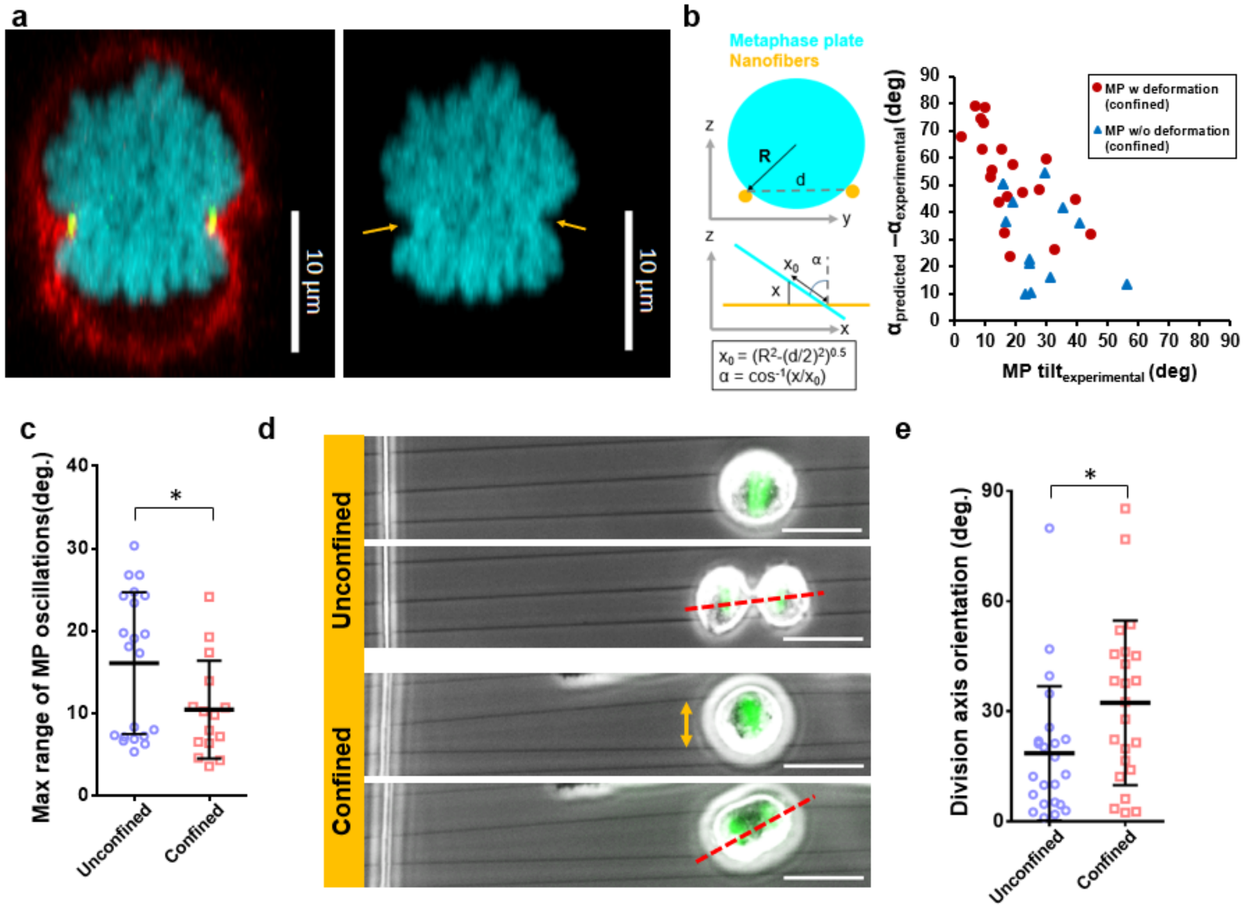


**Supplementary Figure S6: Effect of mechanical confinement on 3D morphology and orientation of the metaphase plate** a) Representative YZ cross-sectional images of confined cell showing constraining fibers hold metaphase plate in position and can also induce deformations locally (actin (red) and DNA Histone H2B (cyan), b) Positive difference between theoretical and experimentally measured tilt angle of MP shows that in the deformed case (red circles) the tilts are lower, thus a larger difference between predicted and measured values, c) Comparison of max range of MP oscillations between confined and unconfined cells demonstrating mechanical stability imparted by fiber-based confinement reduces MP movements, d, e) Representative images and statistical comparison of MP tilt angle for unconfined and confined category cells

Appendix

### Nanonet force microscopy

We used our previously reported ‘Nanonet force microscopy’ to quantify cell-forces at various stages of the cell cycle. We employ nanofiber networks consisting of large diameter (2 µm) support fibers placed ~350 µm apart and an orthogonal layer of small diameter (250 nm) fiber layer with an inter-fiber spacing of ~ 10 µm. The fiber layers are fused at their junctions though solvent vapor exposure. This leads to generation of fixed-fixed boundary conditions at both ends of the small diameter fiber. Fibers are modeled as Euler-Bernoulli beams. Cellular force exertion on fibers is dependent on the stage of the cell-cycle.

During interphase, since focal adhesion organization demonstrate distinct clustering at the cell extremities, the force exertion on each fiber can be considered as a 2-point load. In this stage, cells exert both horizontal and vertical forces on the fibers. The direction of this resultant force exertion is taken along the average orientation of the actin stress fibers ($\alpha_{SF}$) emerging from each focal adhesion cluster. During mitotic entry, breakdown of actin stress fibers occurs, and contractile forces (inward fiber deflection) are generated by the actin-based retraction fibers and their orientation ($\alpha_{RF}$) determine the direction of resultant force exertion. Orientation of the actin stress fibers and retraction fibers are quantified from immunofluorescent staining of actin.

During mitosis, cells adopt rounded shape and push fibers outward. In this configuration, the force exertion can be estimated by using a vertical force model, with the expansive forces from the stiff actin cortex acting orthogonal (α=90°) to the fibers.

| **Stage in cell cycle** | **Cytoskeletal/ adhesion organization** | **Force-bearing structure** | **Direction of cell forces** |
|---|---|---|---|
| *Interphase* | 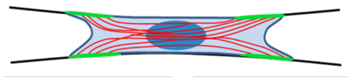 | *Actin stress fibers* | 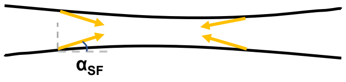  |
| *Mitotic entry* | 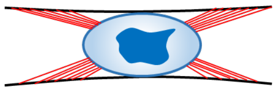 | *Actin-based retraction fibers* | 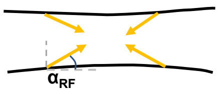  |
| *Mitosis* | 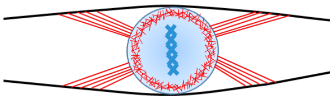 | *Actin cortex* | 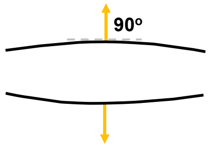  |

Characterization of forces at different stages of the cell-cycle based on cytoskeletal organization

For force quantification, we quantify the cell-mediated fiber deflections and these deflection profiles of fibers bonded to the support fibers on either end, serve as input for the optimization framework to calculate the cell forces. Specifically, the ‘*load updating for* *finite element models*' method has been utilized and the optimization to determine cell forces, is performed through a MATLAB Gradient Based Optimizer (GBO).

The formulation for the cell forces have been reported previously by our group. Briefly each taut fiber can be estimated as a beam with an axial force. Finite Element Method (FEM) can be used discretize the beam into a finite number of uniform straight beam elements as shown in this schematic. Shear forces and moments at two nodes are shown by $Y_1, M_1, Y_2$ and $M_2$.

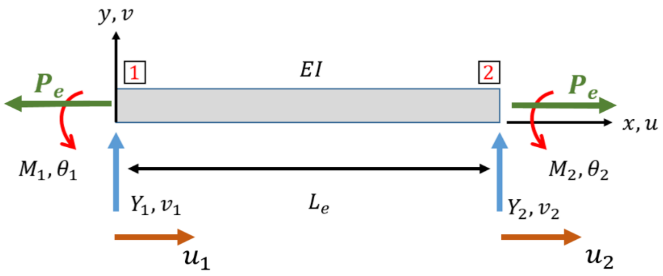


Finite beam element with axial force ($P_e$), shear force (Y) and Bending moment (M)

Using Castigliano's theorem to model the strain energy of each beam element, we can derive the interrelationship between the shear forces and moments exerted on the beam elements and the corresponding displacements.

$$\{\boldsymbol{F}\} = [[\boldsymbol{k}] + [\boldsymbol{n}]]\{\boldsymbol{q}\}$$

Here $[\boldsymbol{k}]$ and $[\boldsymbol{n}]$ will be referred to as the basic stiffness matrix and incremental stiffness matrix or geometric stiffness matrix respectively.

Here

$$\{\boldsymbol{F}\} = \begin{Bmatrix} Y_1 \\ M_1 \\ Y_2 \\ M_2 \end{Bmatrix}, \quad [\boldsymbol{k}] = \frac{EI}{Le}\begin{bmatrix} \frac{12}{Le^2} & \frac{6}{Le} & -\frac{12}{Le^2} & \frac{6}{Le} \\ \frac{6}{Le} & 4 & -\frac{6}{Le} & 2 \\ -\frac{12}{Le^2} & -\frac{6}{Le} & \frac{12}{Le^2} & -\frac{6}{Le} \\ \frac{6}{Le} & 2 & -\frac{6}{Le} & 4 \end{bmatrix}, \quad [\boldsymbol{n}] = \frac{P_e}{10}\begin{bmatrix} \frac{12}{Le} & 1 & -\frac{12}{Le} & 1 \\ 1 & \frac{4Le}{3} & -1 & -\frac{Le}{3} \\ -\frac{12}{Le} & -1 & \frac{12}{Le} & -1 \\ 1 & -\frac{Le}{3} & -1 & \frac{4Le}{3} \end{bmatrix}, \quad \{\boldsymbol{q}\} = \begin{Bmatrix} v_1 \\ \theta_1 \\ v_2 \\ \theta_2 \end{Bmatrix}$$

$P_e$ is the axial tensile force in the element and $L_e$ is the length of the element.

The optimization framework is based on minimizing the following objective function

$$g(x) = \frac{1}{2}\left|\left|\boldsymbol{V_{EXP}} - \boldsymbol{V_{FEM}}\right|\right|^2$$

Here $\boldsymbol{V_{EXP}}$ represents the vector of vertical displacements generated from interpolation of the experimental vertical displacements and $\boldsymbol{V_{FEM}}$ is the vector of computational vertical displacements from the finite element model.

The overall flowchart for the optimization routine is as follows:

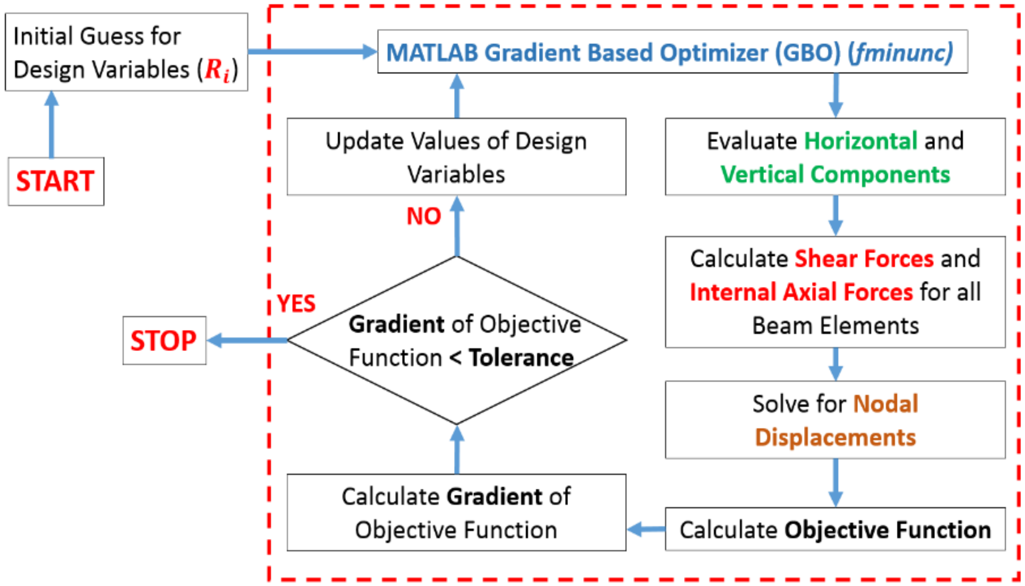


The tolerance is set at $10^{-10}$ and once the optimization criteria is reached, the cell forces corresponding to a particular fiber deflection profile is characterized.

The following properties of the beam are used for cell force calculations(1, 2).

| Property | Value |
|---|---|
| Length (µm) | 350 |
| Diameter (nm) | 250 |
| Young's Modulus (GPa) | 0.97 |
| Pre-Tension (nN) | 201.2 |


1. K. Sheets, J. Wang, W. Zhao, R. Kapania, A. S. Nain, Nanonet Force Microscopy for Measuring Cell Forces. *Biophys. J.* **111**, 197–207 (2016).

2. B. Tu-Sekine, *et al.*, Inositol polyphosphate multikinase is a metformin target that regulates cell migration. *FASEB J.* **33**, 14137–14146 (2019).

**Supplementary Movie S1**: HeLa cell expressing Histone H2B GFP dividing on a single fiber (timestamp-h:min) and scale bar 20μm.

**Supplementary Movie S2**: HeLa cell expressing Histone H2B GFP dividing on a orthogonal arrangement of fibers (timestamp-h:min) and scale bar 20μm.

**Supplementary Movie S3**: HeLa cell expressing Histone H2B GFP dividing on a fiber doublet (timestamp-h:min) and scale bar 20μm.

**Supplementary Movie S4**: HeLa cell expressing Histone H2B GFP dividing on multiple fibers (timestamp-h:min) and scale bar 20μm.

**Supplementary Movie S5**: HeLa cell expressing Histone H2B GFP dividing on flat glass coverslips (timestamp-h:min) and scale bar 20μm.

**Supplementary Movie S6:** Metaphase plate (MP) oscillations over the course of mitosis for HeLa cell dividing on a single fiber. Red arrow is included in the GFP channel for visual tracking of the MP oscillations (timestamp-h:min) and scale bar 20μm.

**Supplementary Movie S7:** Metaphase plate (MP) oscillations for HeLa cell dividing on crossing fibers (timestamp-h:min) and scale bar 20μm.

**Supplementary Movie S8:** Metaphase plate (MP) oscillations for HeLa cell dividing on fiber doublets (timestamp-h:min) and scale bar 20μm.

**Supplementary Movie S9:** Reduced level of metaphase plate (MP) oscillations for HeLa cell dividing on multiple fibers (timestamp-h:min) and scale bar 20μm.

**Supplementary Movie S10:** High-speed recording of division event of Hela cell on single fibers, showing movements of the rounded cell during metaphase (timestamp-h:min:sec) and scale bar 20μm.

**Supplementary Movie S11:** High-speed recording of HeLa division on crossing fibers (timestamp-h:min:sec) and scale bar 20μm.

**Supplementary Movie S12:** High-speed recording of HeLa division on a fiber doublet (timestamp-h:min:sec) and scale bar 20μm.

**Supplementary Movie S13:** High-speed recording of HeLa division on multiple fibers (timestamp-h:min:sec) and scale bar 20μm.

**Supplementary Movie S14:** Hela cell expressing Histone H2B GFP, entrapped in fiber doublets deflecting fibers outward during mitotic rounding (timestamp-h:min) and scale bar 20μm.

**Supplementary Movie S15:** 3-dimensional rendering of a confocal z-stack showing a rounded HeLa cell in metaphase, mechanically confined within a fiber doublet. Actin: red, Fibers: green and chromosomes: blue

**Supplementary Movie S16:** HeLa cell dividing under confinement, showing tilted orientation of division angle (timestamp-h:min) and scale bar 20μm.
